# Biological Synthesis of Silver Nanoparticles with Aqueous Extract of *Azadirachta indica* Leaf and Its Antimicrobial Activity on Uropathogenic MDR and ESBL Producing *Escherichia coli*

**DOI:** 10.64898/2026.04.10.716351

**Authors:** Payel Das

## Abstract

**Background:** The plethora of *Escherichia coli (E. coli)* strains that show multidrug resistance (MDR) has risen. The production of extended-spectrum beta-lactamases (ESBL) enzymes has greatly aided in this. There have been speculations on the effectiveness of silver nanoparticles in treating drug-resistant infections.

**Aim:** This research aims to utilize *Azadirachta indica* (neem) leaf extract for the production of silver nanoparticles (AI-AgNPs), considering its medicinal and antimicrobial properties. Additionally, this research intends to evaluate the efficacy of these nanoparticles on resistant *E. coli* infections.

**Method:** The antimicrobial activities and cytotoxicity of silver nanoparticles synthesized using neem leaves extract were tested on MDR and ESBL producing *E. coli* strains, as well as on Human Embryonic Kidney-293 (HEK-239) cell line, respectively. The characterization of silver nanoparticles was done by UV-Vis Spectrophotometry, Fourier-transform infrared spectrometry (FTIR), Dynamic Light Scattering-Zeta Potential (DLS-Zeta), X-Ray Diffraction (XRD), Scanning Electron Microscope (SEM), Transmission Electron Microscope (TEM), and Energy Dispersive X-Ray Spectroscopy (EDAX).

**Results:** The synthesized nanoparticles were spherical, smooth, and stable with an average size of approximately 74 nm. The Minimum inhibitory concentration (MIC) of AI-AgNPs was 9.5 µg/ml, and the Minimum bactericidal concentration (MBC) was 121 µg/ml. The IC50 value for AgNPs was 369 µg/mL for HEK-293 cell line exposure.

**Conclusion:** This study showed that the biosynthesis of silver nanoparticles from *A. indica* extract is highly effective, exhibiting strong antibacterial activity against multidrug-resistant bacteria while exhibiting low toxicity to normal human kidney cells. Hence, biosynthesized silver nanoparticles may be useful as antimicrobial materials for infection control because of their remarkable antibacterial activity.

## 1. Introduction

Silver was employed by the Romans, Persians, Egyptians, and Greeks approximately 5000 years ago for food storage. (1). Silver’s antimicrobial properties are likely why it has been used for millennia to make eating and drinking utensils. (2). One of the most active research fields nowadays appears to be nanotechnology (3, 4) and AgNPs are more effective than other nanoparticles for biological purposes, particularly when used against microorganisms and insects (5). Prospective applications of AgNPs include wound healing, implantations, wound cleaning, sensing, catalysis, covering other surfaces, and more (6). The exact process by which AgNPs function is still unknown. Certain research suggests that AgNPs can kill antibacterial cells by deactivating proteins, damaging cell membranes, and inducing thiol reactions leading to enzyme inactivation and cell death (7-9). DNA damage, inflammation, and oxidative stress may also contribute (10).

A variety of techniques, including chemical, biological, and physical techniques, have been employed to create AgNPs (11). Traditional AgNPs synthesis involved toxic chemicals, prompting the adoption of eco-friendly “Green synthesis.” This method, preferred for its environmental friendliness, cost-effectiveness, scalability, and avoidance of high-pressure, high-temperature, energy, and toxic substances, offers a simpler alternative to conventional approaches (12). Because of their antioxidant qualities and ability to reduce metal compounds in their respective nanoparticles, bacteria, fungi, and plants have been used in a considerable number of studies on the green synthesis of AgNPs (2). The best capping material for stabilizing AgNPs is produced by plant extracts (13). Some findings demonstrated that coated AgNPs are less hazardous than uncoated AgNPs, and toxicity depends on the nature of the coating (14). Compared to silver ions (Ag^+^) and other coating materials, AgNPs coated with plant material are less toxic (15). Concerns have been raised about the potential toxicity of using AgNPs inside the human body (16-18). Smaller nanoparticles exhibit potent antimicrobial effects but higher cytotoxicity. Therefore, AgNPs are often combined with other substances to enhance effectiveness and mitigate toxicity (6).

Neem (*A. indica*), a widespread plant in India and neighbouring regions, belongs to the Meliaceae family and is renowned for its medicinal properties (19). Its phytochemicals, primarily terpenoids and flavanones, serve as both reducing and capping agents, aiding in nanoparticle stabilization (20). Nanoparticles synthesized with neem extract as a capping agent demonstrate heightened antibacterial activity (11).

People living in impoverished and developing nations are more susceptible to contracting microbiological infections because of poor residential cleanliness and inadequate infrastructure (21). A prevalent nosocomial illness in low- and middle-income countries is uropathogenic *E. coli* infection with a high AMR burden (22, 23). The occurrence of MDR among *E. coli* isolates was elevated, with ESBL production identified as a leading factor contributing to MDR (24). Clinical isolates are commonly linked to infections in humans or animals and tend to be more virulent because of the selective pressure present within the host organism (25). These isolates are often exposed to antibiotics, which contributes to their increased resistance to antimicrobial treatments compared to environmental bacteria (26). Additionally, they may carry multiple resistance mechanisms and exhibit different behaviours than bacteria found in natural environments (27). Considering the advantages of green synthesis, medicinal properties of neem as well as its antimicrobial activity, this study aims at the synthesis of AgNPs using aqueous neem leaves extract (AI-AgNPs) and to evaluate its efficacy in ameliorating infection caused by resistant clinical strains of *E. coli*. Some articles show how *E. coli* infections are impacted by silver nanoparticles made with *A. indica* (2, 11). This study uniquely demonstrates the green synthesis of stable silver nanoparticles using *A. indica* leaf extract and their potent activity against clinically isolated MDR and ESBL-producing *E. coli*. Unlike previous works, it highlights the efficacy of neem-mediated AgNPs on resistant uropathogenic strains with minimal cytotoxicity, offering an eco-friendly and biocompatible alternative for antimicrobial applications.

## 2. Methodology

### 2.1. Collection and identification of *A. indica* leaves

Fresh leaves were collected from the trees harvested in the locality of Kolkata, West Bengal, India. Collected leaves were washed thoroughly with Milli-Q water and dried in air. Genomic DNA was isolated using the GenElute Plant Genomic DNA Miniprep Kit (Sigma). The presence of 18S rRNA unique to *A. indica* validated the species of the plant. PCR was performed to identify 18S rRNA using primer sequence (5’-3’) Forward: GCAGAATCCCGTGAACCATC and Reverse: GCTTGTTCTCACCACCGATC. The PCR conditions for the amplifications were as follows: 95°C for 2 min followed by 40 cycles of 95°C for 1 min, 58°C for 30 sec, and 72°C for 1 min, with a final extension at 72°C for 10 min. The amplified PCR products were detected by Agarose gel electrophoresis (AGE) and visualized using GelDoc to capture the digital image for analysis.

### 2.2. Preparation of leaf extracts from *A. indica* (Neem) leaves

Fresh green leaves were collected from the neem tree, washed with Milli-Q water, and dried in air until the water evaporated completely. They were finely chopped and weighed. 20 grams of these finely chopped leaves were then boiled with 100 ml of Milli-Q water for 10 minutes, cooled, and filtered using Whatman filter paper Grade 1 (Pore size: 11µm). The obtained extract was then stored at 4°C.

### 2.3. Preparation of Silver Nanoparticle (AI-AgNPs)

Silver nitrate (AgNO3) (Sigma) was employed to produce a 100 ml solution of aqueous AgNO3 with a concentration of 10^-2^ M. Subsequently, varying concentrations (2%, 4%, 6%, 8%, and 10%) of neem extract were individually added to 1 ml of the 10-2 M AgNO3 solution, and the total volume was adjusted with Milli-Q water to dilute the AgNO3 to 10-3 M. This mixture was then subjected to incubation at room temperature. The alteration in colour from colourless to brown serves as a confirmation of the formation of AgNPs. The nanoparticles were gathered through centrifugation, followed by washing the pellet three times with deionised water to eliminate any undesired substances. The solution containing AgNPs derived from *A. indica* was subjected to centrifugation at speeds of 10000 rpm, 12000 rpm, and 15000 rpm for a duration of 10 minutes (Fig. 1).

**Figure 1:**
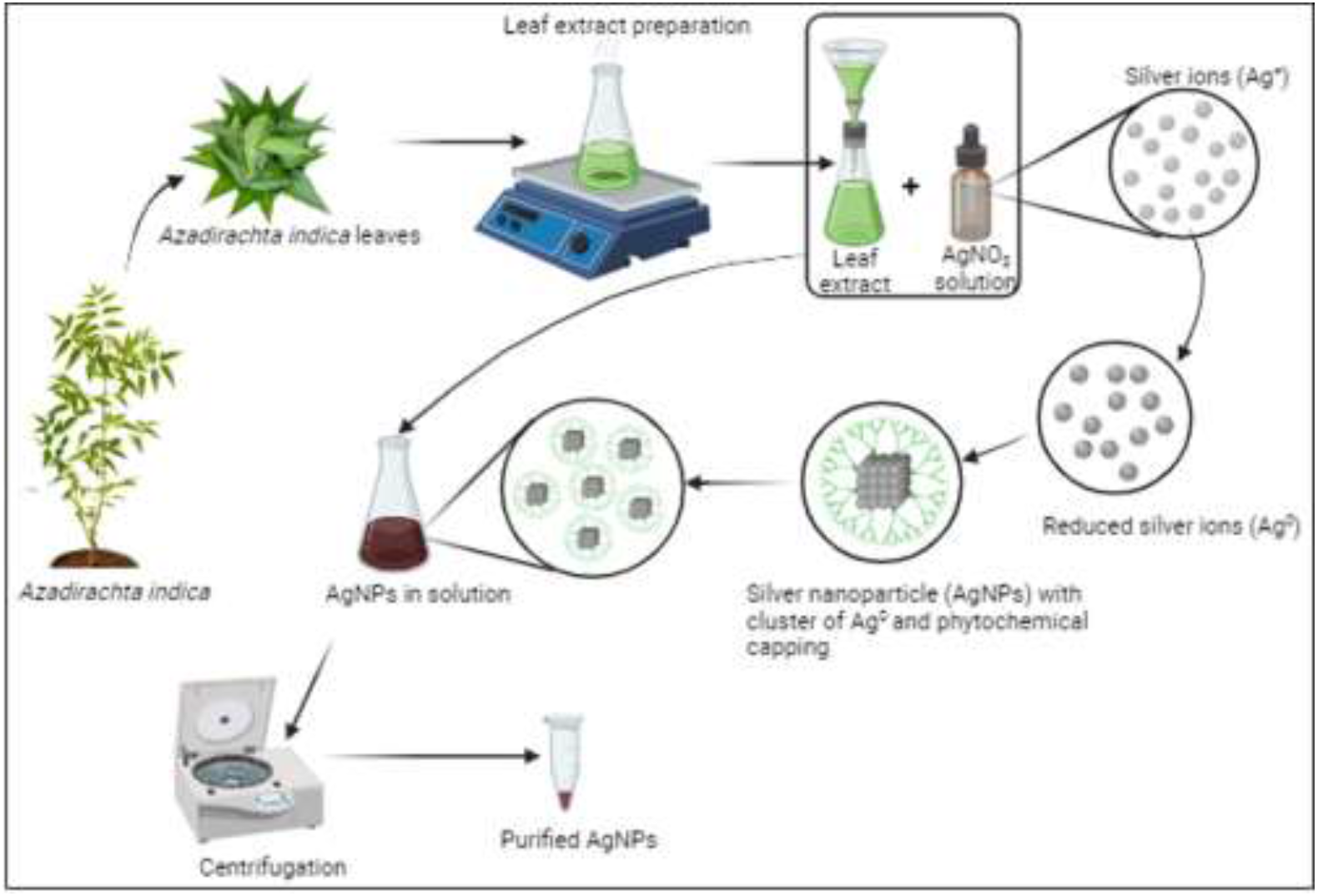
Schematic diagram of green synthesis of silver nanoparticles.

### 2.4. Characterisation of silver nanoparticles

### 2.4.1. UV-Visible spectrum analysis

Absorbance of the AgNPs has been taken at 24 hours, 48 hours, 72 hours, and 96 hours, respectively. Synthesis of AgNPs was measured by a UV-visible spectrophotometer (Systronics) at a wavelength difference of 10nm from 300 to 600 nm.

#### 2.4.2. FTIR analysis

A Fourier Transform Infrared Spectrometer (FTIR) (JASCO FT/IR-6300) was used to analyse AgNPs at a resolution of 4 cm^-1^ and in the 4000–400 cm^-1^ range to detect various functional groups that are responsible for capping and stabilising the particles.

#### 2.4.3. X-ray Diffraction Analysis

Using an X-ray diffraction (XRD) spectrometer (Bruker, Advance D8) equipped with Cu Kα1 radiation (λ= 1.54060 Å) in the 2θ range of 10° to 90°, the mean crystal size, phase composition, and other structural details of AgNPs were verified. The device was operated at 30 kV with a current of 30 mA.

#### 2.4.4. DLS & Zeta Potential analysis

AgNPs were measured for hydrodynamic size and zeta potential using the Malvern Panalytical Zetasizer Advanced Series (Zetasizer Pro).

#### 2.4.5. SEM and EDAX analysis

Scanning electron microscopy (SEM) (Zeiss Evo 18 Special Edition, Germany) has been done to check their morphology using platinum-coated drop-casted AgNPs onto the cover glass. Sizes of the AgNPs were measured using Image J software. Energy-dispersive X-ray (EDAX) analysis has been done for elemental composition using the same instrument with dried AgNPs powder.

#### 2.4.6. TEM analysis

The size, dispersion, and morphology of the nanoparticles have been examined using Transmission Electron Microscopy (TEM). A drop of nanoparticle was applied to a copper TEM grid coated with carbon to prepare the sample, which was then dried in a vacuum chamber for 45 minutes. TEM was performed using JEOL JEM-2100 HR, operated at an accelerating voltage of 200 kV and 0.23 nm resolution. Sizes of the AgNPs were measured using Image J software.

### 2.5. Bacterial culture preparation

88 pre-cultured clinical samples were obtained from different UTI patients at different hospitals and private laboratories located in West Bengal. They were phenotypically and genotypically confirmed as MDR and ESBL. Bacterial cultures were drawn in Mueller-Hinton broth (MHB) and incubated overnight at 37°C. Standard suspensions of these cultures are prepared by matching a specific turbidity standard (0.5 McFarland standard).

*E. Coli* (ATCC 35218), *K. pneumoniae* (ATCC 700603), *E. coli* (ATCC 25922), and *K. pneumoniae* (ATCC 25955) were used as reference strains for comparison.

### 2.6. Antimicrobial assay

To detect the antimicrobial activity of these AI-AgNPs, MIC and MBC were performed.

#### 2.6.1. MIC Determination

MIC is the minimum concentration at which an antimicrobial agent can inhibit the growth of a bacterium. To assess the MIC, 800µg of AgNPs are dissolved in 1 ml of sterile distilled deionised water in a sterile Eppendorf tube and mixed by vortexing to form a homogenised solution. AgNPs are serially diluted from 1-10 numbered wells in 96 96-well plate. Then 100 µl of bacterial culture was added to it to attain a concentration of 5×10^5^ CFU/ml in each well. MHB and Meropenem were used as negative and positive controls. It is then incubated for 18-24 hours at 37°C. After incubation, 40 µl of Iodonitrotetrazolium (INT) was added to each well, and a change of colour was observed. Formation of colour indicated the growth of bacteria, and no change in colour indicated that bacteria could not grow.

The same procedure is followed to determine the antimicrobial activity of *A. indica* leaf extract. In this case, serial dilution starts with 50% of the leaf extract.

#### 2.6.2. Determination of MBC

For determination of MBC, 10 µl from the wells that did not show any change in colour were again inoculated on a sterile 96-well plate containing 190µL of MHB and further incubated for 18-24 hours. After incubation, 50 µl of INT was added to each well, and a change of colour was observed. A change in colour indicated the presence of live bacterial cells.

All antimicrobial assays were performed in triplicate.

### 2.7. Cytotoxicity study on normal cell line by MTT assay

The cytotoxicity potential of the AgNPs on the human normal kidney cell line (HEK-293) was evaluated by the MTT assay. The HEK-293 cell line was maintained in Dulbecco’s Modified Eagle Medium (DMEM, GIBCO) as per the standard protocol. The active HEK-293 cells (5×10^4^ cells/well) were loaded in 96-well plates and incubated for 1 day at 37 °C. Various concentrations (25-500 μg/mL) of the AI-AgNPs were administered to the growing HEK-293 cells and incubated at 37 °C for 24 h. At the end of the treatment, the medium was added with MTT (0.05 mg/mL) and incubated at 37 °C for 4 h in a CO2 incubator. Then, the plate was centrifuged at 1000 RPM for 10 minutes, the medium was aspirated, and the cells were rinsed with Phosphate Buffered Saline (PBS) and then, the cells were dissolved with 100 μL of Dimethyl Sulfoxide (DMSO), coloured with formazan stain, and blended well. The control represented by DMEM without any treatments. Well, with DMEM without cells and treatment was considered as blank. The microplate reader (iMark Microplate Absorbance Reader, Bio-Rad) was used to read the 96-well plates at 595 nm. The percentage of cell viability was calculated using the following formula:

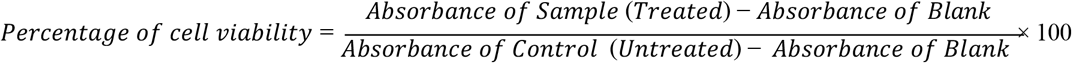

## 3. Results

### 3.1. Identification of *A. indica* leaves

Primers for the *A. indica-*specific 18S rRNA were able to amplify a specific part of the genomic DNA extracted from the collected leaves. The prominent band at 230 bp in AGE confirms *A. indica* (Fig. 2).

**Figure 2:**
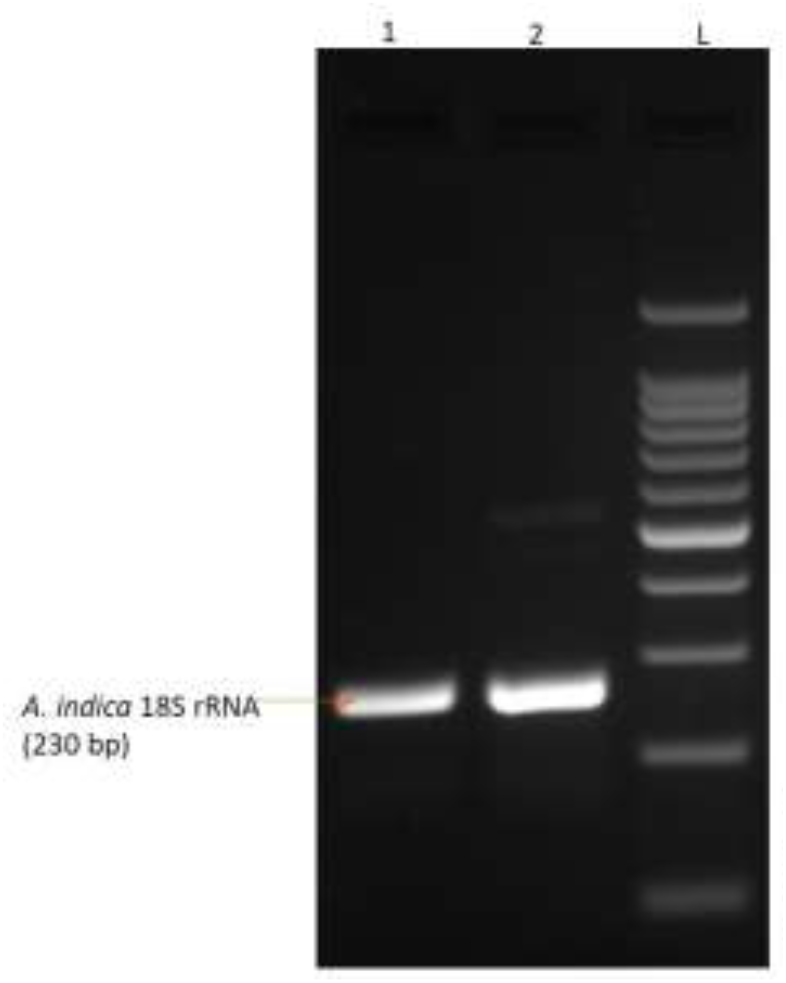
Gel picture of amplified product of *A. indica* 18S rRNA. 1-2: Neem leaf samples; L Ladder.

### 3.2. UV-Visible spectrum analysis

A noticeable shift in colour was observed upon the incorporation of aqueous neem extract into the AgNO3 solution. As shown in the figures, the solution’s colour altered from pale yellow (Fig. 3A) to brown (Fig. 3B). In the 300–600 nm range, UV–vis spectroscopy confirmed the synthesis of AgNPs. In the case of AgNPs from *A. indica* shows a highly intense resonance peak around 440 nm, having a concentration of 8% of plant extract. But a less intense peak had also been found at 380nm in the case of AgNPs with 8% and 6% of plant extract. In the case of 2% and 4% of plant extract, a single intense resonance peak around 440nm had been observed. No such peak was observed in the case of 10% plant extract (Fig. 4).

**Figure 3:**
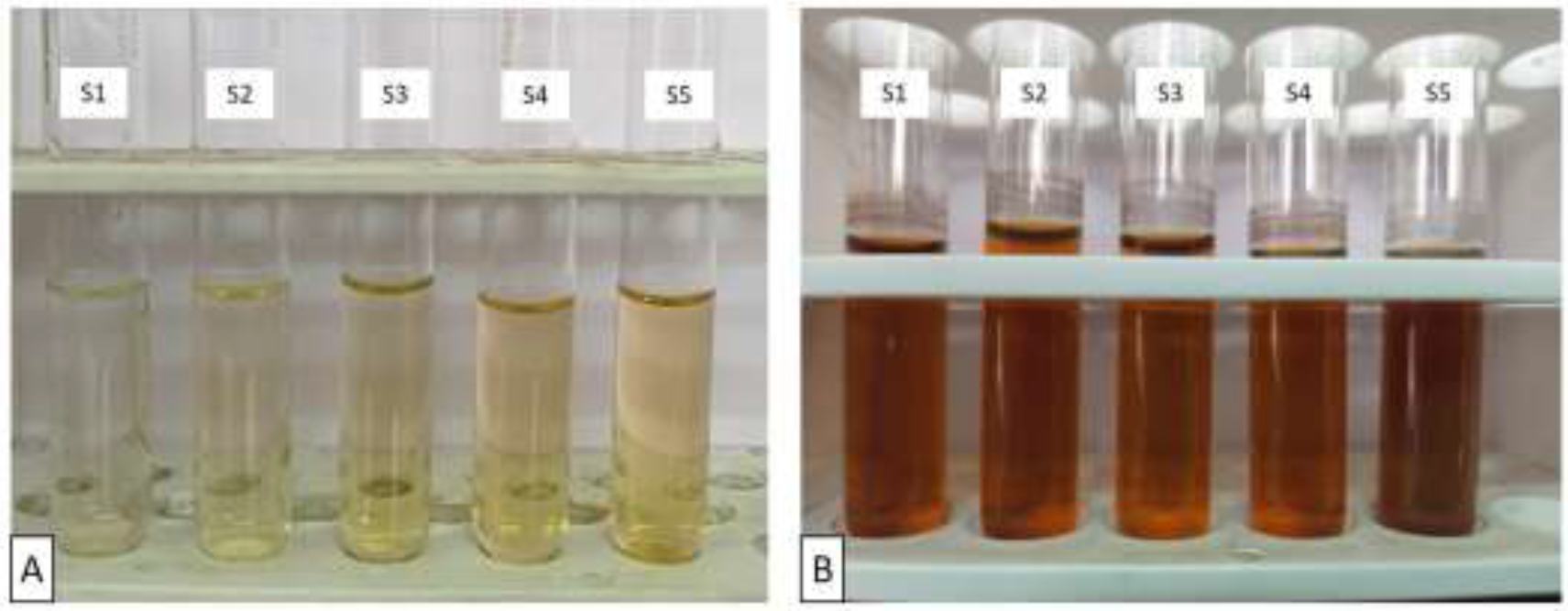
Change in colour of the mixture of AgNO3 and Neem leaf extract (Sl=2% neem leaf extract, S2=4% neem leaf extract, S3=6% neem leaf extract, S4=8% neem leaf extract, S5=10% neem leaf extract). A: Colour development at 0 hour of incubation; B: Colour development after 72 hours of incubation.

**Figure 4:**
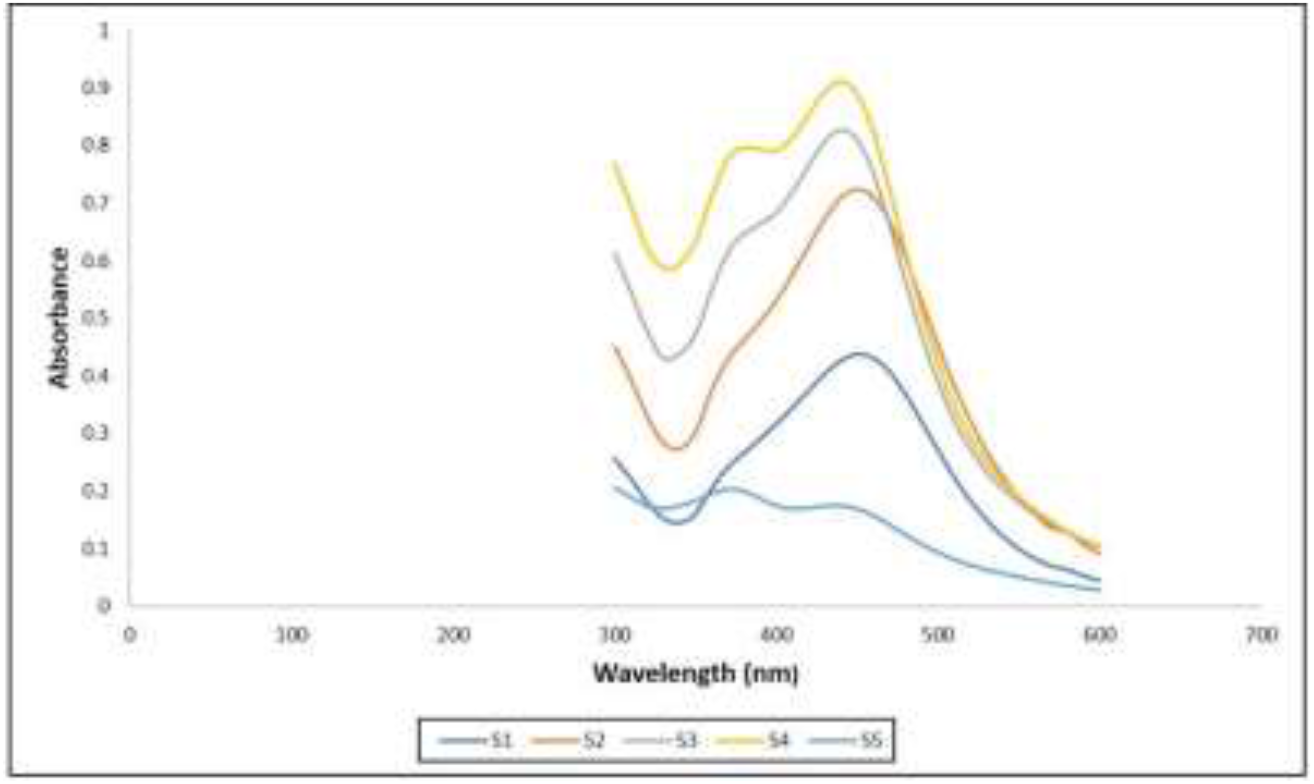
Absorption spectra of silver nanoparticles at various concentrations of Neem leaf extract. S1=2% neem leaf extract, S2=4% neem leaf extract. S3=6% neem leaf extract, S4=8% neem leaf extract, S5=10% neem leaf extract.

The intensity of the resonance peak increased with time (Fig. 5A). After 72 hours of incubation, the increase of this peak intensity became very slight or nearly constant (Fig. 5B).

**Figure 5:**
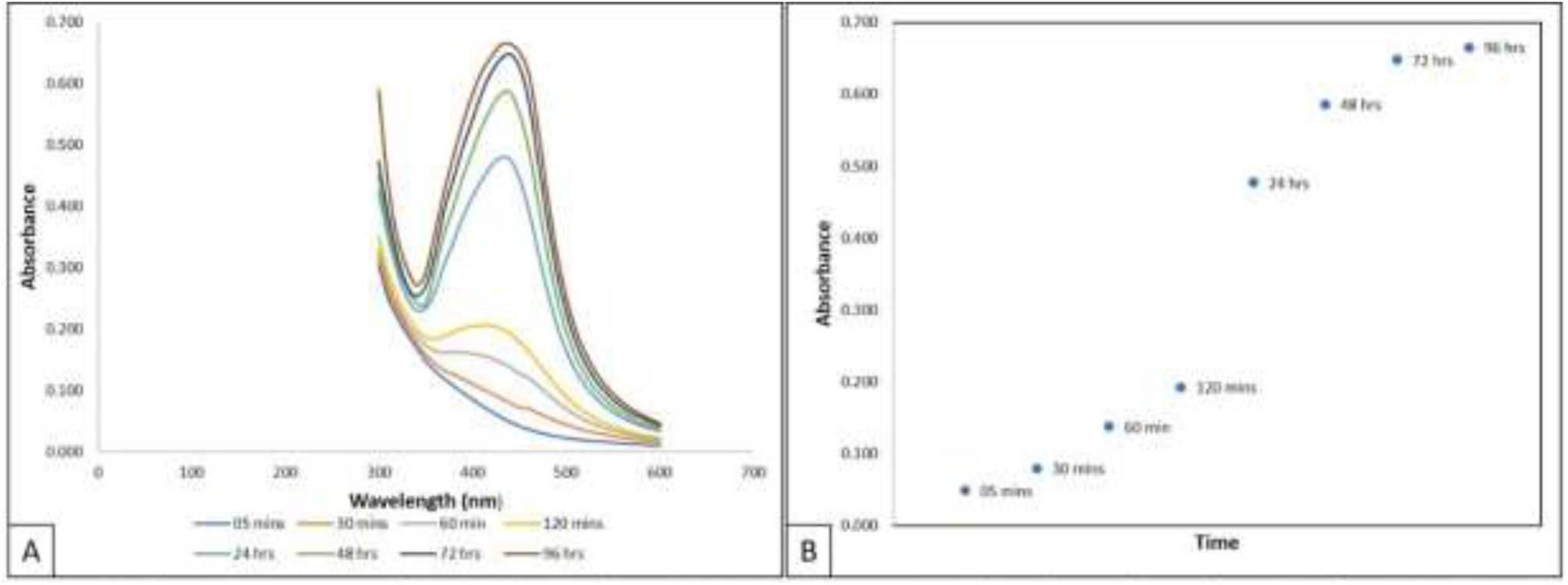
Absorption spectra of *Azadirachta indica* synthesized silver nanoparticles observed as a function of reaction time. A: Absorption spectra of silver nanoparticles observed at eight different reaction times. B: Increase of absorption intensity as a function of reaction time.

The maximum AgNPs pellet was collected by centrifugation at 15000 rpm for 10 minutes at 20°C (Fig. 6).

**Figure 6:**
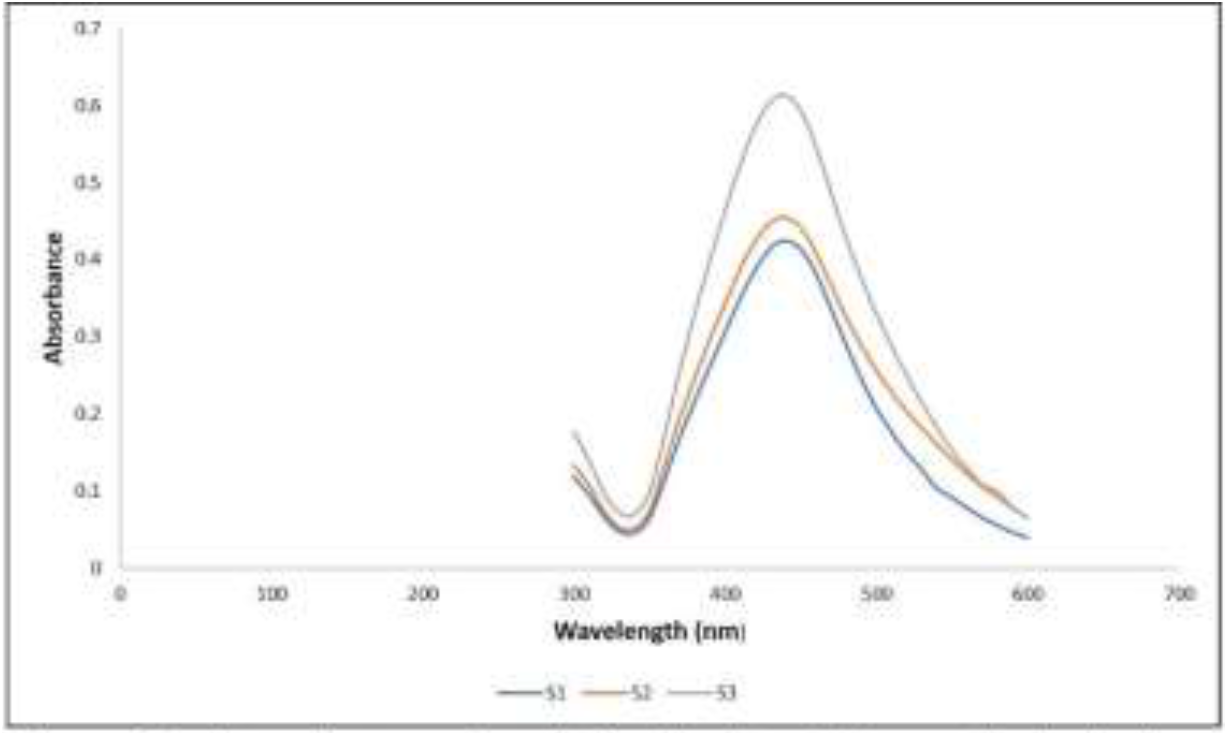
Absorption spectra of *Azadirachta indica* synthesized silver nanoparticles centrifuged at three different speeds. Sl= Centrifuged at 10000 rpm; S2= Centrifuged at 12000 rpm; S3= Centrifuged at 15000 rpm.

### 3.3. FTIR analysis

The active functional groups stabilising AgNPs produced with *A. indica* aqueous leaf extract were determined by FTIR analysis. Fig. 7 displays multiple distinct and prominently apparent peaks throughout the observation range of 400 cm^-1^ to 4000 cm^-1^ (Table 1). Pure leaf extract and AgNPs from leaf extract show a peak at 3429 cm^-1,^ which corresponds to the OH and amine groups. Vibrations at 2925 cm^-1^, 1653 cm^-1^, and 949 cm^-1^ are observed due to C-H, C=C, and =C-H of the alkene, respectively. 2854 cm^-1^ indicates O-H of carboxylic acid and 1749 cm^-1^ corresponds to the C=O of carbonyl group. 1484 cm^-1^ shows vibrations due to the C-C stretch of aromatic ring, and 1181 cm^-1^ indicates the presence of C-N of amines. The presence of a unique geminal dimethyl group corresponds to the peak at 1383 cm^-1,^ and the peak at 1097 cm^-1^ indicates the presence of an ether linkage.

**Table 1:**
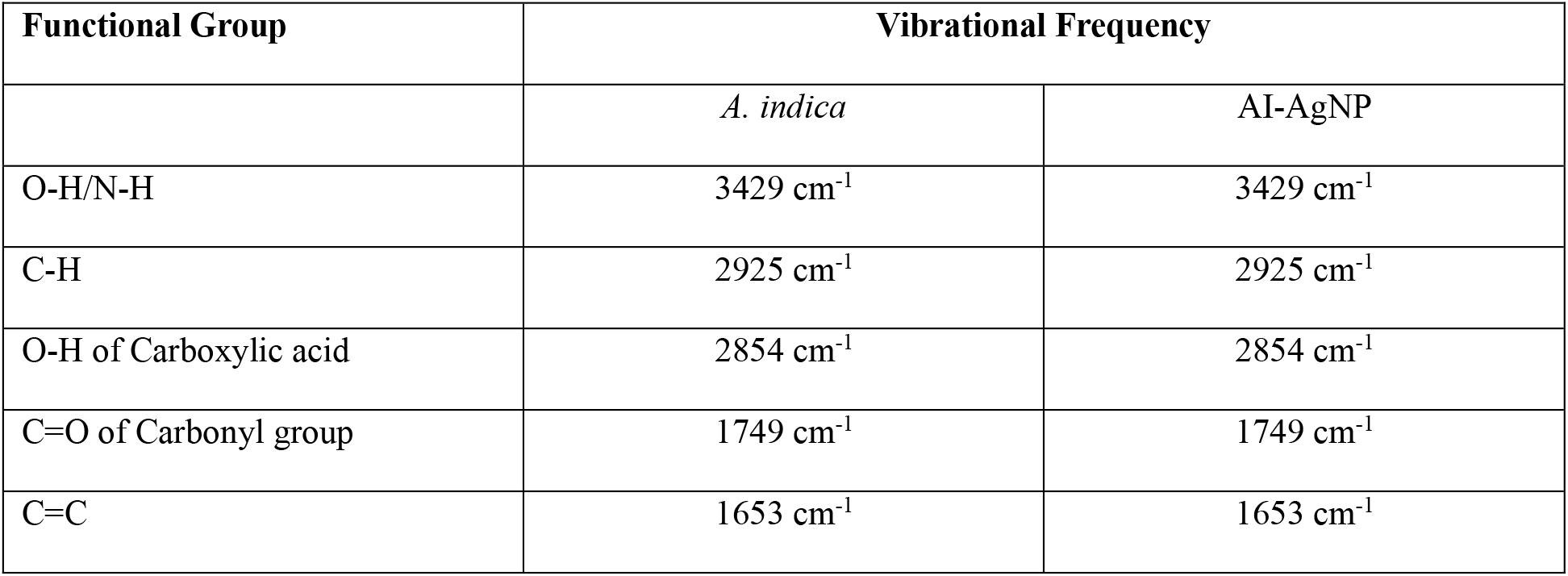
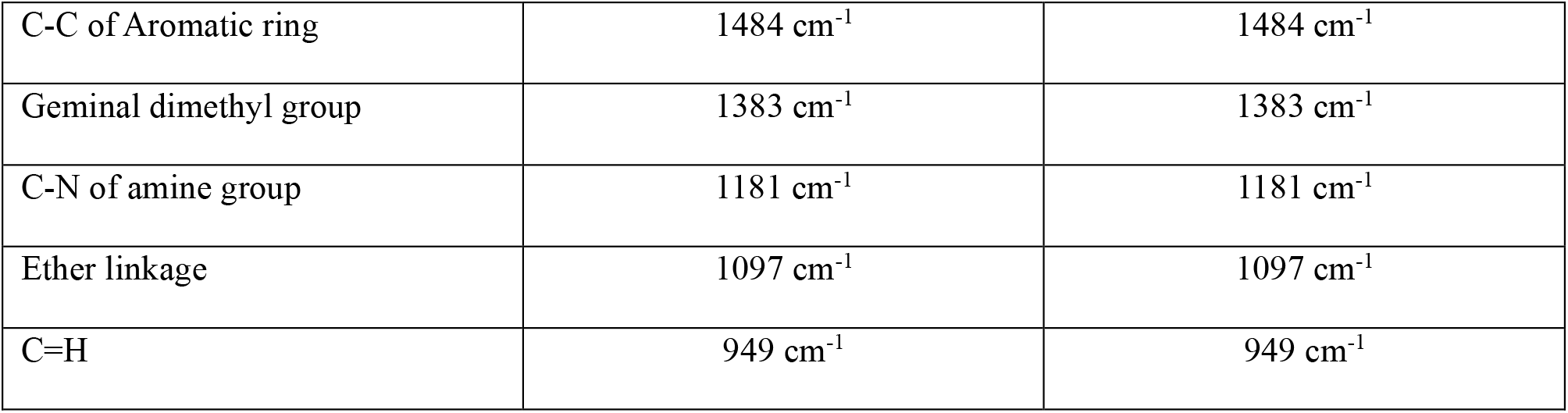
FTIR spectral properties of *A. indica* (Neem) leaf extract and synthesized silver nanoparticles (AI-AgNP).

| Functional Group | Vibrational Frequency |  |
| --- | --- | --- |
|  | *A. indica* | AI-AgNP |
| O-H/N-H | $3429\text{ cm}^{-1}$ | $3429\text{ cm}^{-1}$ |
| C-H | $2925\text{ cm}^{-1}$ | $2925\text{ cm}^{-1}$ |
| O-H of Carboxylic acid | $2854\text{ cm}^{-1}$ | $2854\text{ cm}^{-1}$ |
| C=O of Carbonyl group | $1749\text{ cm}^{-1}$ | $1749\text{ cm}^{-1}$ |
| C=C | $1653\text{ cm}^{-1}$ | $1653\text{ cm}^{-1}$ |
| C-C of Aromatic ring | 1484 cm<sup>-1</sup> | 1484 cm<sup>-1</sup> |
| Geminal dimethyl group | 1383 cm<sup>-1</sup> | 1383 cm<sup>-1</sup> |
| C-N of amine group | 1181 cm<sup>-1</sup> | 1181 cm<sup>-1</sup> |
| Ether linkage | 1097 cm<sup>-1</sup> | 1097 cm<sup>-1</sup> |
| C=H | 949 cm<sup>-1</sup> | 949 cm<sup>-1</sup> |

**Figure 7:**
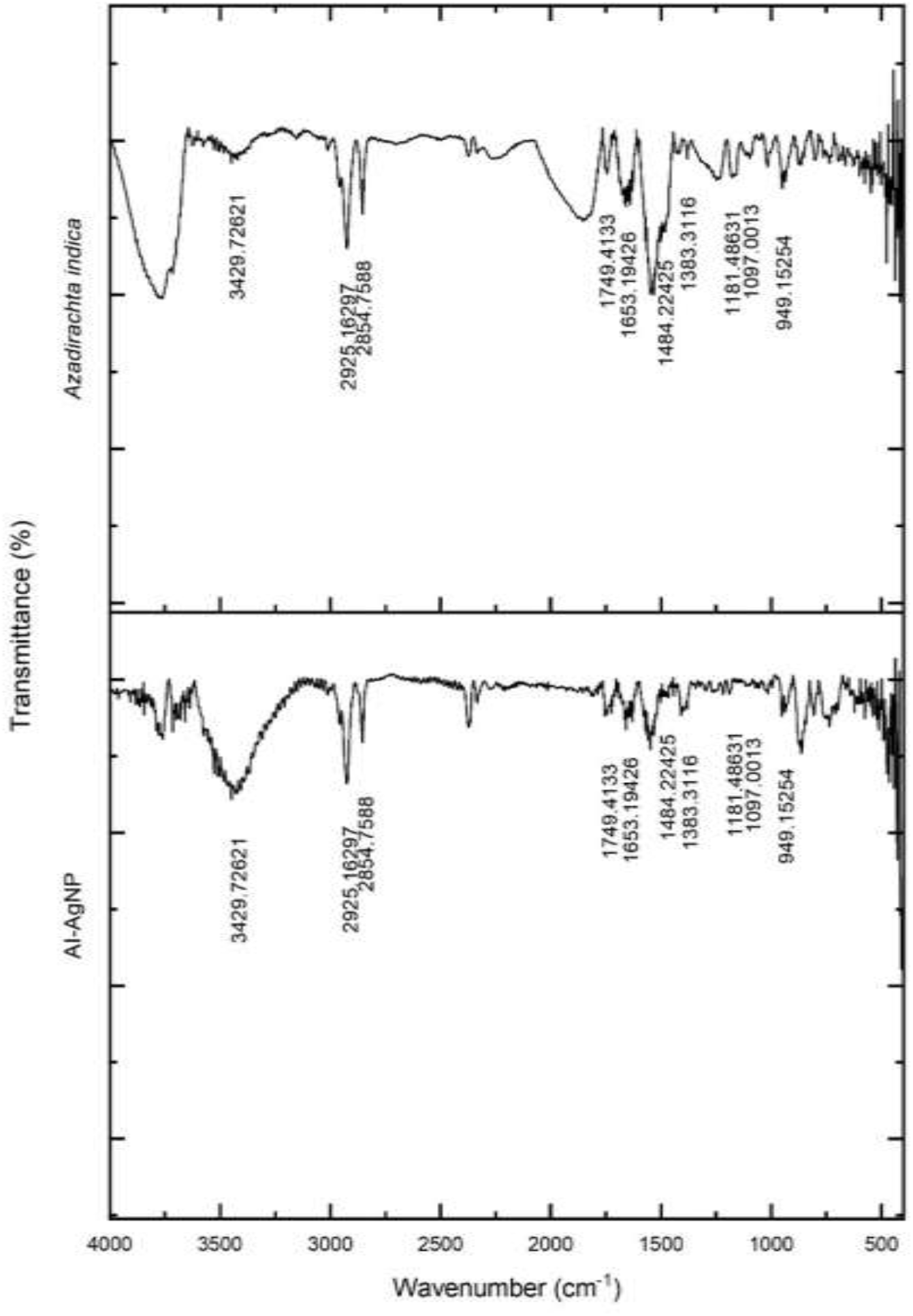
FTIR spectrum of *Azadirachta indica* (Neem) leaf extract and synthesized silver nanoparticles.

### 3.4. XRD Analysis

Diffraction peaks in the XRD pattern are seen at about 2θ positions for the following degrees: 27.864, 32.299, 38.214, 44.351, 46.239, 54.782, 57.542, 64.327, 67.394, 74.753,76.844,77.152, 81.249, and 85.646 (Fig. 8). Joint Committee on Powder Diffraction Standards (JCPDS) file No. 04-0783 lists five of these peaks for pure silver crystals, which correspond to (hkl) values (111), (200), (220), (311), and (222) planes, and have 2θ values of 38.214, 44.351, 64.327, 77.152, and 81.249 degrees in the experimental diffractogram. These peaks are believed to be caused by silver metal. The silver metal’s crystalline structure, which has a face-centred cubic (FCC) symmetry, is compatible with every reflection plane. As expected, the sample displays the strong (111) reflection that is characteristic of FCC materials. Peaks at 27.829, 32.239, 46.226, 54.808, 57.464, 67.427, 74.432, 76.709, and 85.644 with corresponding hkl values of 111, 200, 220, 311, 222, 400, 331, 420, and 422 reflect the presence of chlorargyrite crystals. Reflection planes for the chlorargyrite crystal were compared with the standard chlorargyrite crystals (JCPDS file no. 31-1238).

**Figure 8:**
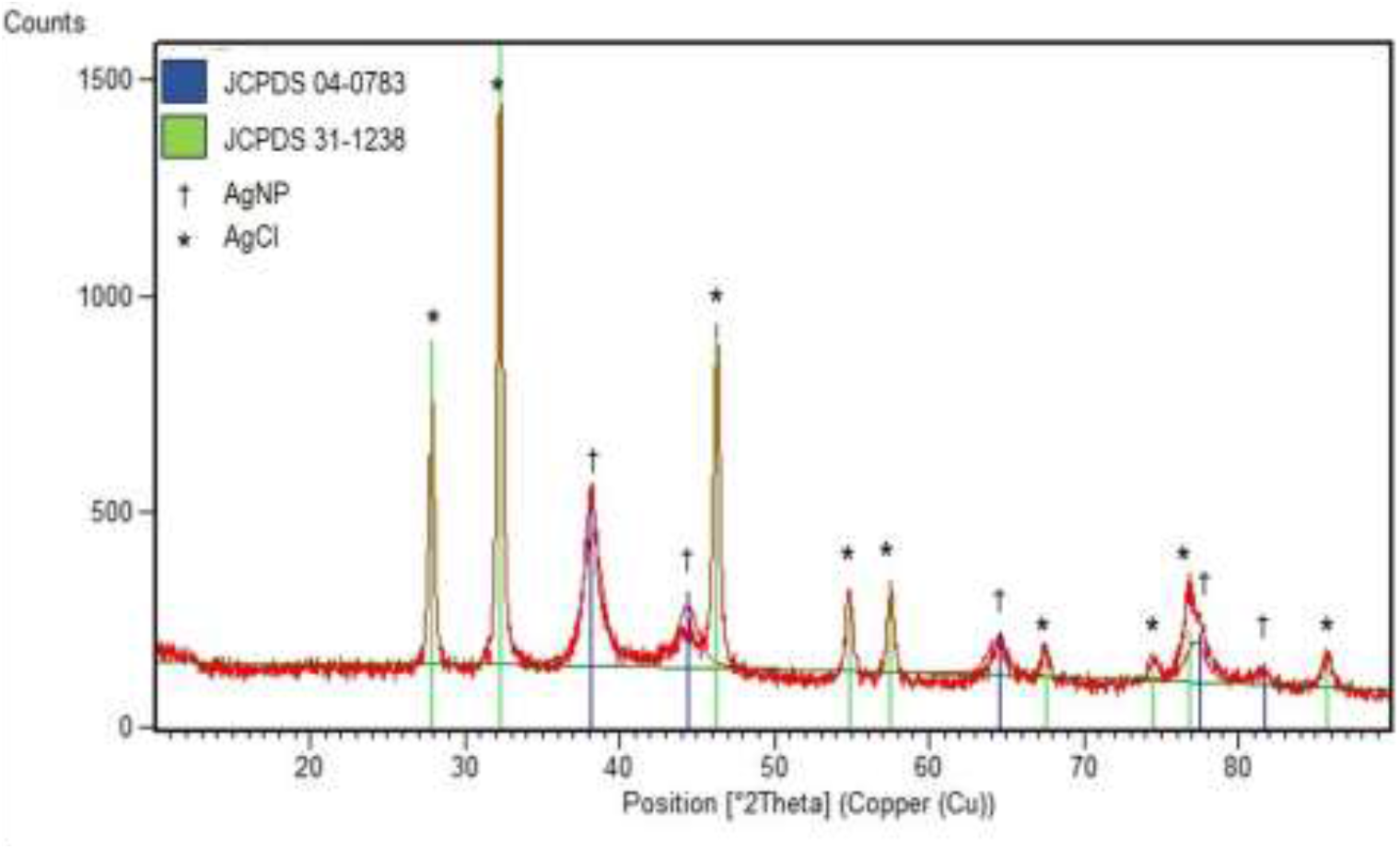
XRD pattern of *Azadirachta indica* synthesized silver nanoparticles.

The crystallite size (D) of AgNPs was determined using the Debye−Scherrer equation (Table 2):

**Table 2.** XRD results of *A. indica* synthesized AgNPs.

| SL No. | Peak Position | hkl | Cosθ | Sinθ | FWHM (°) | β (radians) | Crystallite Size (nm) | Av. Crystallite Size (nm) | Interplanar Spacing (nm) | Lattice Constant (nm) | Av. Lattice Constant (nm) |
| --- | --- | --- | --- | --- | --- | --- | --- | --- | --- | --- | --- |
| 1 | 38.2140 | 111 | 0.9449 | 0.3273 | 0.1968 | 0.0034 | 42.7208 | 73.6674 | 0.2353 | 0.4076 | 0.4089 |
| 2 | 44.3510 | 200 | 0.9260 | 0.3774 | 0.08872 | 0.0015 | 96.6960 |  | 0.2041 | 0.4082 |  |
| 3 | 64.3270 | 220 | 0.8465 | 0.5323 | 0.15744 | 0.0027 | 59.6078 |  | 0.1447 | 0.4093 |  |
| 4 | 77.1520 | 311 | 0.7818 | 0.6236 | 0.144 | 0.0025 | 70.5680 |  | 0.1235 | 0.4097 |  |
| 5 | 81.2490 | 222 | 0.7590 | 0.6511 | 0.106 | 0.0019 | 98.7445 |  | 0.1183 | 0.4098 |  |

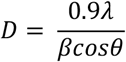

Where the Cu Kα X-ray wavelength λ = 0.15406 nm, θ is Bragg’s diffraction angle (° or radian), and β (radian) is the full width at half-maximum (FWHM) of the peak. The average crystallite size was found to be approximately 73.6674 nm.

The interplanar spacing (d) was calculated using Bragg’s formula (Table 2):

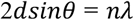

Where n is the order of the diffraction pattern. In the present case, n is equal to 1.

The Lattice constant (a) had been estimated using the formula (Table 2):

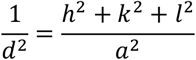

The estimated average lattice constant, 0.4089 nm, is quite near to the typical lattice parameter of AgNPs, which is about 0.4086 nm (JCPDS file number 04-0783).

### 3.5. DLS analysis

The size and distribution profile (Fig. 9) of the small particles in suspension were ascertained by employing the dynamic light scattering (DLS) technique. Synthesised AgNPs have a polydispersity index (PDI) value of 0.368, according to the DLS data. The intensity vs. size distribution curve revealed that the particles’ average size was 73.74 nm. AgNPs have a zeta potential of -28.04 mV (Fig. 10).

**Figure 9:**
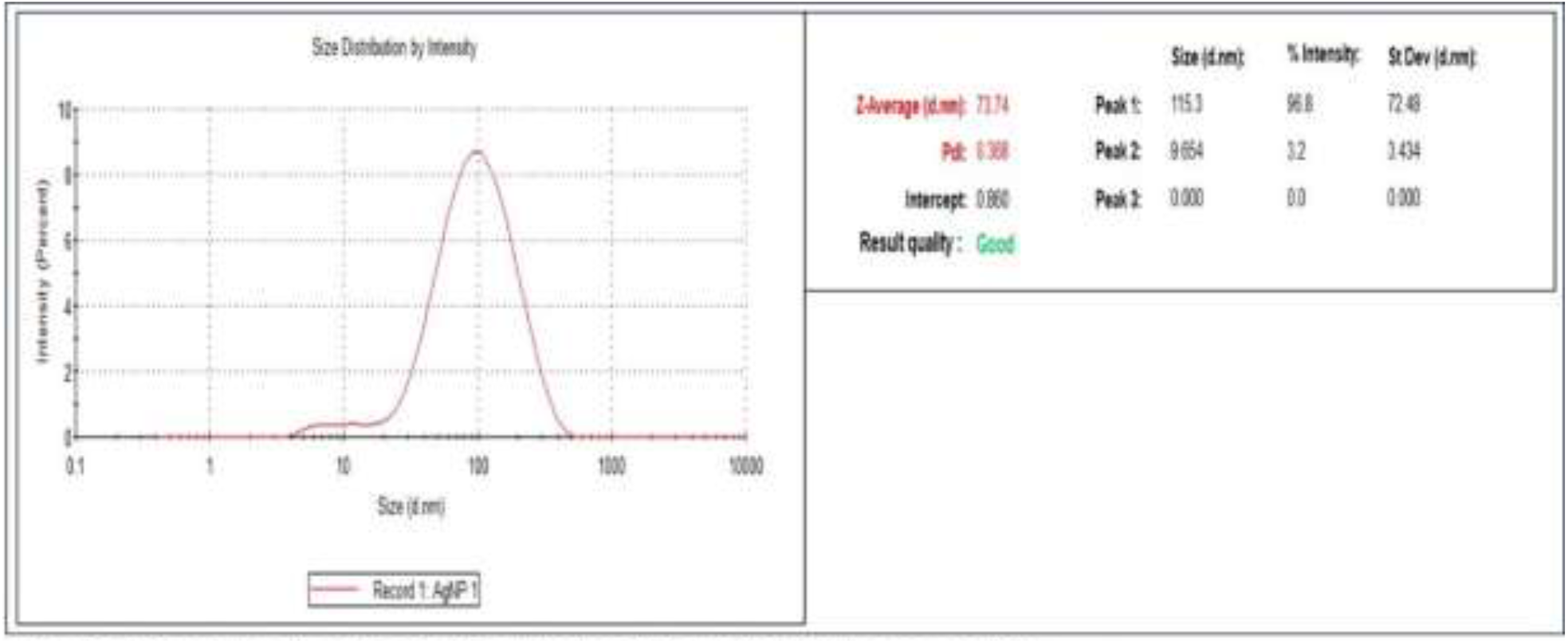
DLS spectrum of particle size of *Azadirachta indica* synthesized silver nanoparticles.

**Figure 10:**
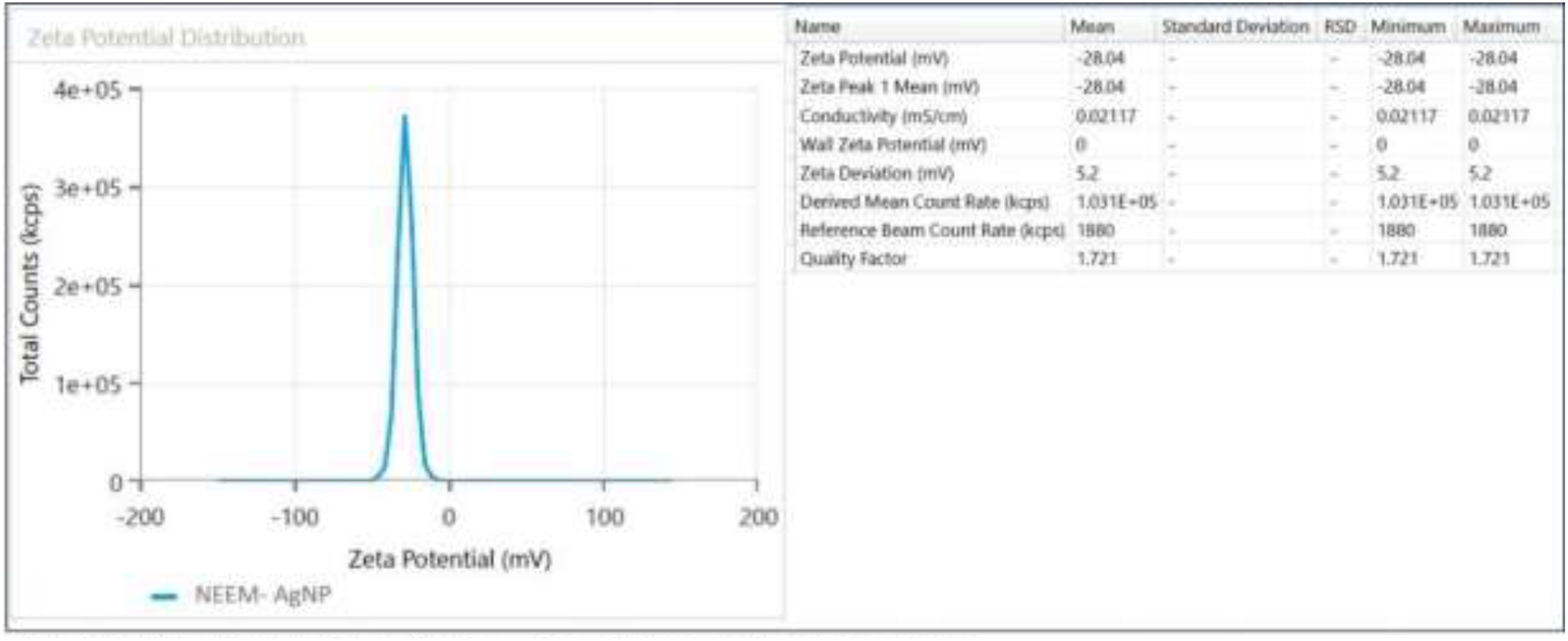
Zeta Potential of *Azadirachta indica* synthesized silver nanoparticles.

### 3.6. SEM and EDAX analysis

**The** SEM image (Fig. 11) represents spherical particles with few clusters. These particles are smooth-surfaced. Sizes of the particles were estimated by measuring their diameter by Image J software. The average particle size was 72.0164±9.803 (Mean±SD). EDAX plot (Fig. 12) of AgNPs shows strong signal from silver (61.3%), chlorine (15.9%), platinum (12.2%), carbon (8.2%), and oxygen (2.3%).

**Figure 11:**
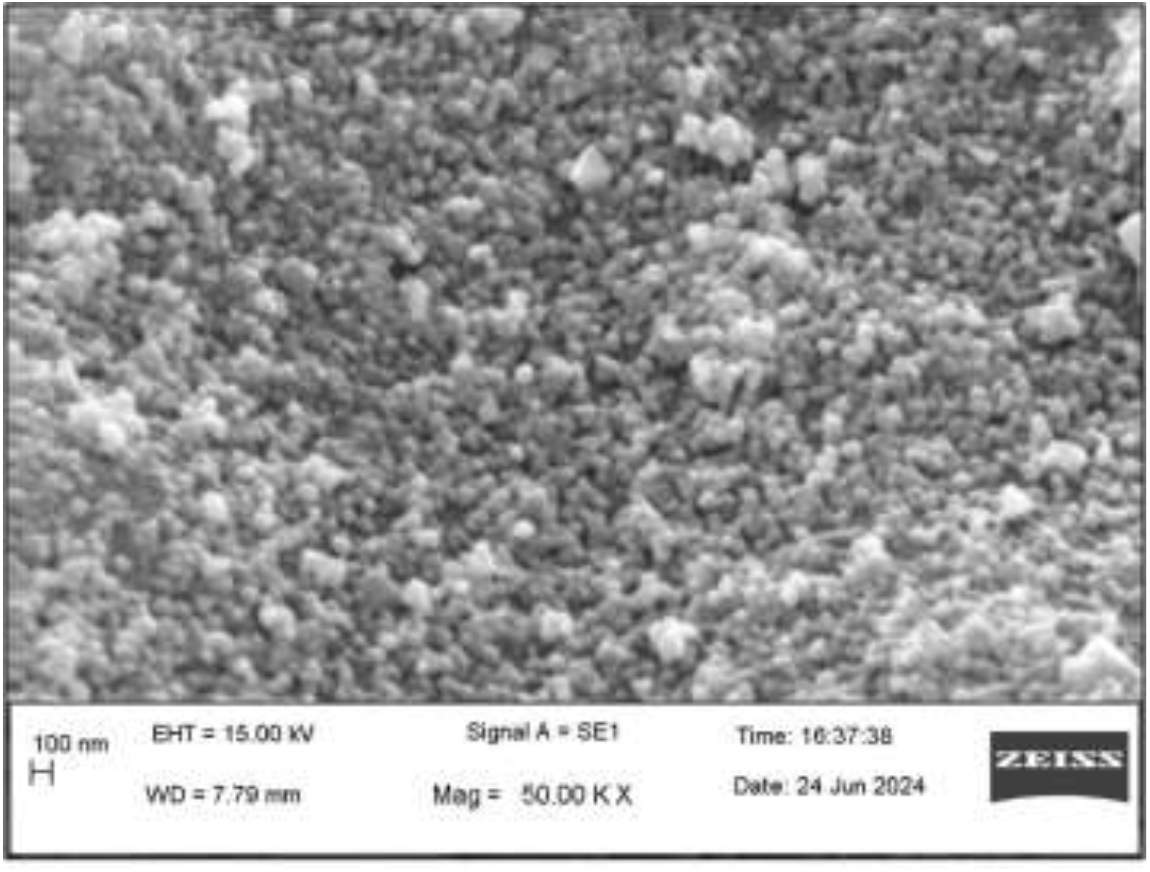
SEM Image of *Azadirachta indica* synthesized silver nanoparticles.

**Figure 12:**
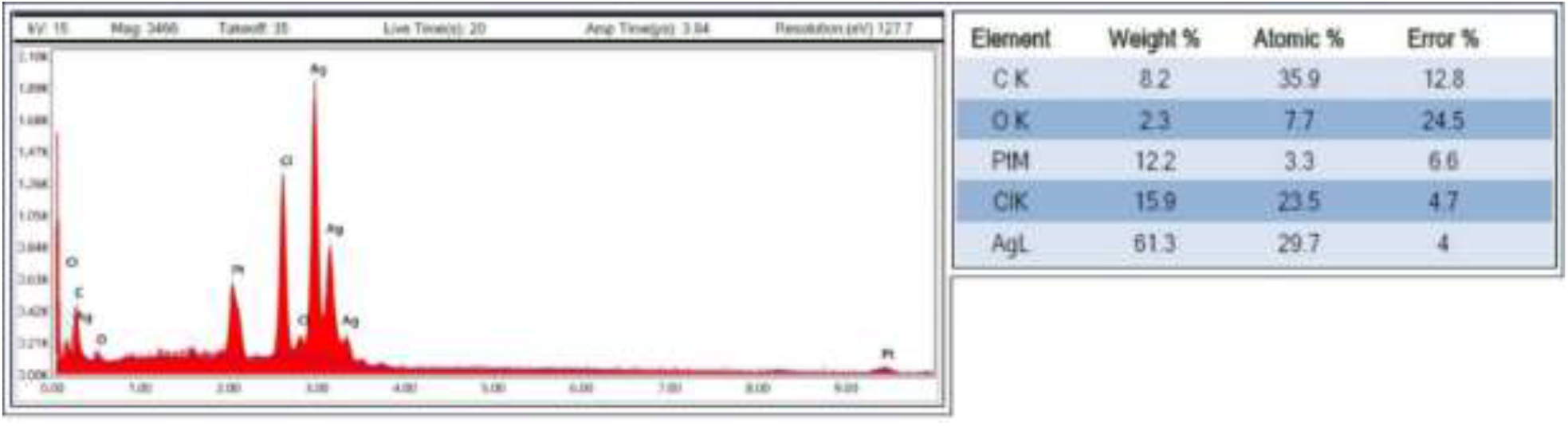
EDAX pattern of *Azadirachta indica* synthesized silver nanoparticles.

#### TEM analysis

TEM image (Fig. 13) clearly shows smooth-surfaced surfaced almost spherical, nanosized particles. In the higher magnification image, silver nanoparticles of 57-98 nm size were observed (Fig. 3A). However, the average size of the nanoparticles was found to be 73.51±13.11 nm.

**Figure 13.**
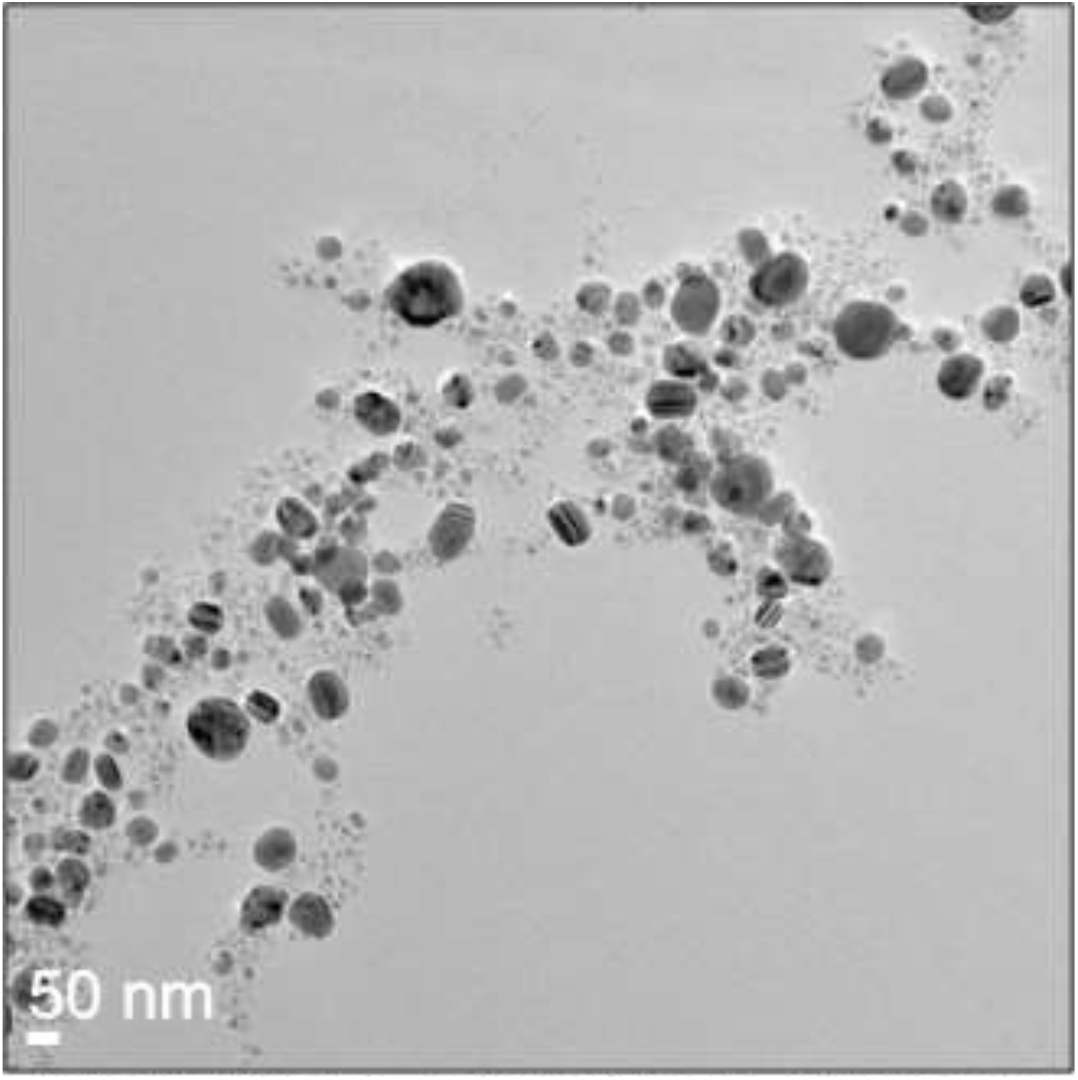
TEM image of *Azadirachta indica* synthesized silver nanoparticles.

### 3.7. Antimicrobial analysis

A microdilution method was used to evaluate the MIC of the green-synthesised AgNPs, with values ranging from 800 to 1.5625 µg/ml. At 12.5 µg/ml of AgNPs, growth inhibition was seen in 33% of the tested bacterial isolates (Fig. 14). There were comparable inhibitory effects observed at doses ranging from 3.125 to 6.25 µg/ml. Neem leaf extract is unable to provide this kind of inhibitory effect on its own (Fig. 15). The concentration that was shown to be most efficient in preventing the growth of MDR and ESBL producing *E. coli* isolates was 9.4993±0.72679 µg/ml (Mean±SE) (95% CI: 8.0547-10.9439). For 35.2% of the bacterial isolates, the AgNPs’ MBC was 200 µg/ml; however, in certain cases, *E. coli* may also be destroyed at doses of 50–100 µg/ml (Fig. 15). These clinical isolates could be eliminated with an average concentration of 121.0227±7.64143 µg/ml (Mean±SE) (95% CI: 105.8345-136.2109).

**Figure 14:**
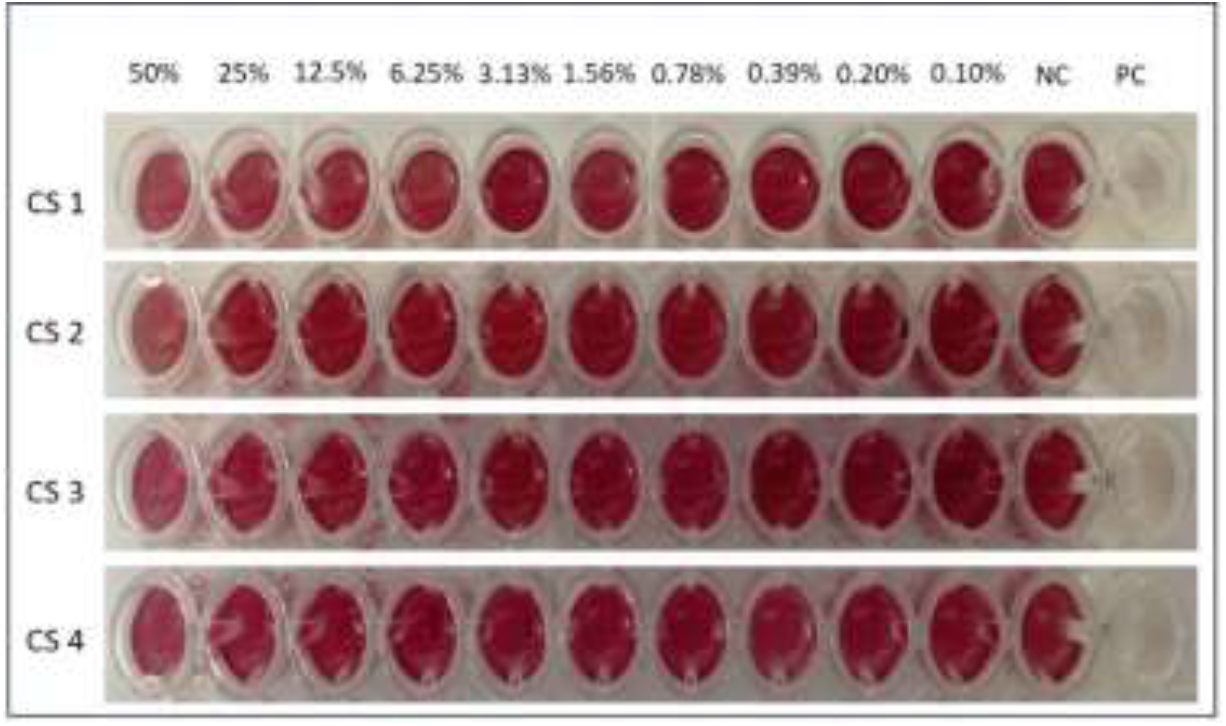
Antimicrobial activity of *Azadirachto indica* leaf extract by MIC Analysis. Development of colour indicates bacterial growth. CS1= Clinical sample 1; CS2= Clinical sample 2; CS3= Clinical sample 3; CS4= Control *E. coli* ATCC 25922; NC= Negative Control; PC= Positive Control.

**Figure 15:**
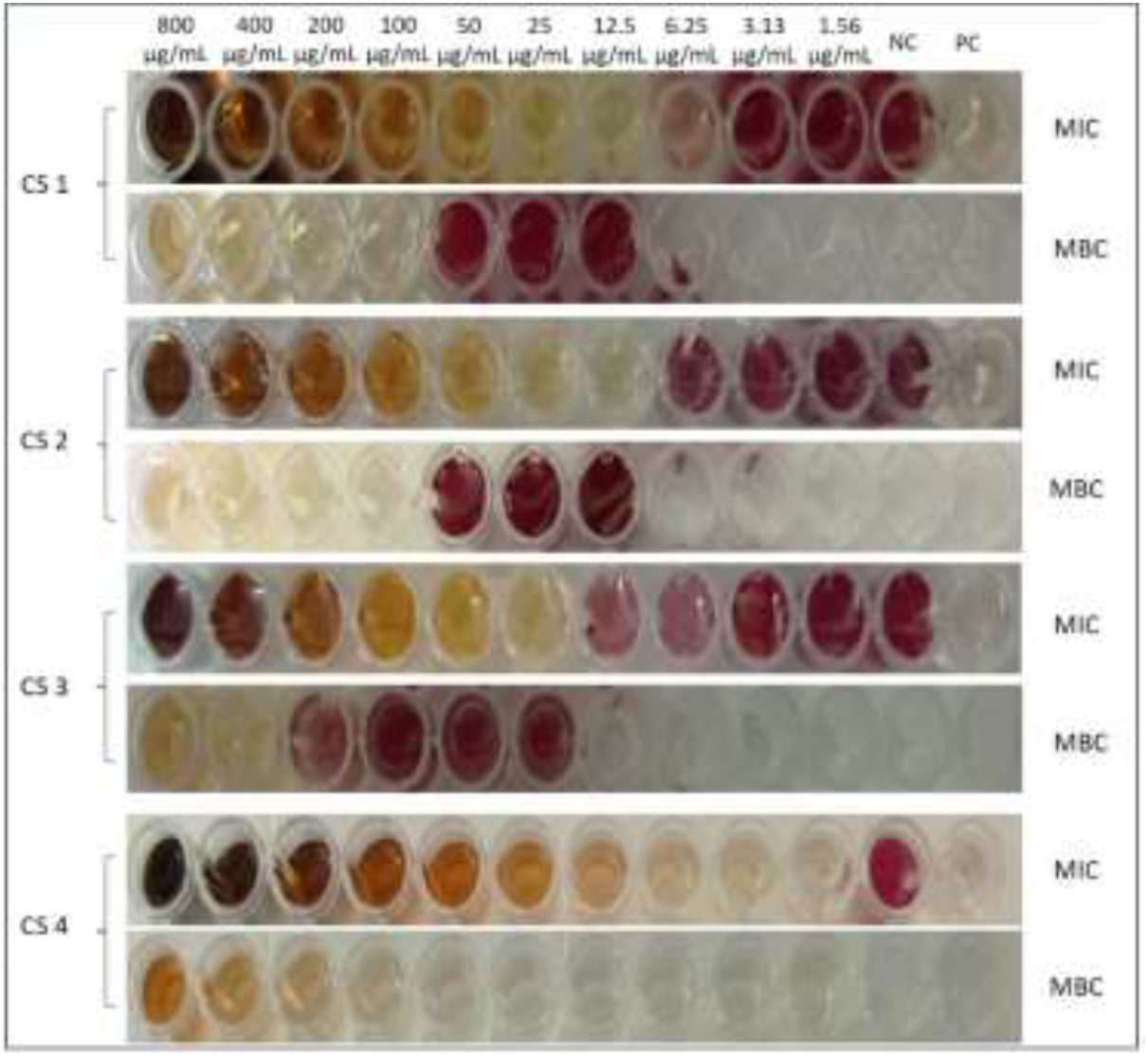
Antimicrobial activity of *Azadirachta indica* synthesized silver nanoparticles by MIC and MBC Analysis. Development of Purple colour indicates bacterial growth. CS1= Clinical sample 1; CS2= Clinical sample 2; CS3-Clinical sample 3; CS4=Control *E. coli* ATCC 25922; NC= Negative Control; PC= Positive Control.

### 3.8. Cytotoxicity study on normal cell line by MTT assay

A decrease in cellular proliferation was observed when HEK 293 was treated with AI-AgNPs in a dose-dependent manner. 50% antiproliferative action (IC50) of AI-AgNPs was exerted against a normal human kidney cell line at a dose of 369 µg/ml after 24 hours of incubation (Fig. 16).

**Figure 16:**
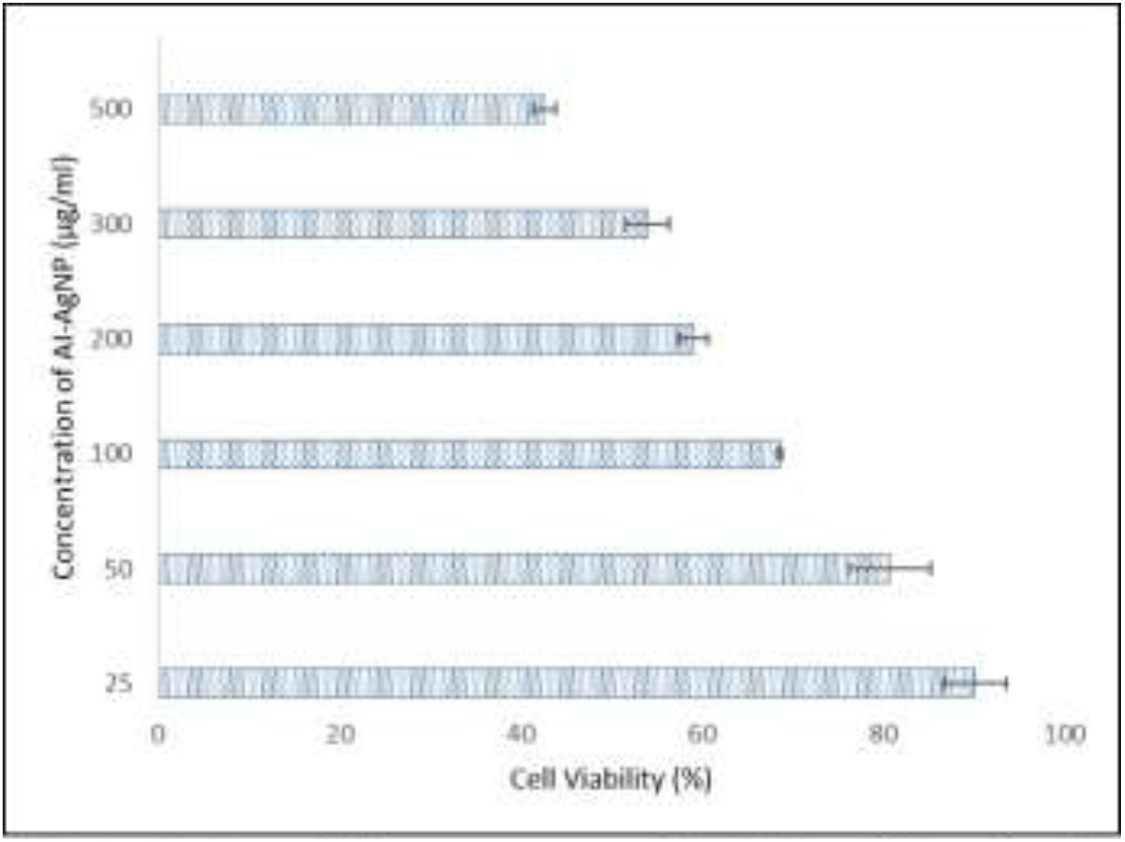
Cytotoxicity measurement of HEK293 cell line after 24 hours of incubation with Al-AgNP. Data are represented as Mean±SD.

## Discussion

AgNPs are well known for their ability to fight off bacteria and have been in medical use for quite some time. In this study, the goal is to create AgNPs using water-based neem leaf extract, which is recognised for its antimicrobial activity. The presence of the 230 bp long PCR product, which was amplified with a primer specific to *A. indica*, demonstrates that the leaves belong to the same species. A colour shift indicated that the metal nanoparticles had been formed by the bio-reduction of aqueous silver ions after being exposed to the broth of boiled *A. indica* leaves. The presence of natural reducing agents such as terpenoids and flavanones in neem leaf extract is what causes the silver salt to be reduced to AgNPs, changing its colour from pale yellow to a deep brown over time. The metal nanoparticles’ surface plasmon oscillations are excited, which results in the formation of this colour (11). The silver surface plasmon resonance (SPR) band is found to occur at 430 nm and to consistently grow in intensity with reaction time without changing its peak wavelength. Several other plant extracts produced comparable outcomes (28, 29). Particle size and the solution’s refractive index affect the SPR band (30). The dipole resonance redshifts as the particle size rises and the field across it becomes nonuniform (31). However, a quadrupole resonance mode is created as the particle size goes over 80 nm, exerting a secondary peak at a lower wavelength (32). In this study, 8% and 6% leaf extract exhibit a secondary and less intense peak at 380nm. The formation of bigger nanoparticles larger than 80 nm could be the cause of this occurrence. At 430 nm, the 4% leaf extract had a more intense peak than the 2% and no secondary peak. Therefore, it was decided to move further using 4% neem leaf extract. AgNPs production depends on the ratios of phytochemicals and AgNO3 (33). 10% leaf extract fails to exert such an SPR band. At high quantities, terpenoids and flavanones might not be able to convert Ag^+^ into AgNPs.

The first SPR band was seen 60 minutes after the neem leaf extract was added to the salt solution. The SPR band’s peak becomes almost constant after 72 hours, meaning that no more silver salt is available for reaction. A maximum amount of AgNPs had been gathered at 15000 rpm as smaller particles were pelleted down with the increase in centrifugal force.

The biomolecules in charge of effectively stabilizing and capping the produced metal nanoparticles were determined by analysing the infrared spectra of the green nanoparticles. The alkene, aromatic, hydroxyl, carbonyl, and amine moieties found in *A. indica* have a potent potential to reduce metal, as demonstrated by the IR spectrum, and thus, offer to cap and prevent AgNPs from clumping (11, 34). Adsorbed flavanones on the surface of metal nanoparticles are suggested by the strong bands of ether linkage at 1097 cm^-1^ (34). Hence, the leaf extract of *A. indica* serves as both a capping and a stabilizing agent.

The XRD pattern reveals that the synthesised AgNPs are crystalline and have face-centred cubic symmetry. Of all the diffraction peaks at 2θ positions of 38.214°, 44.351°, 64.327°, 77.152°, and 81.249° corresponding to (111), (200), (220), (311), and (222) reflection planes. The asterisk-marked peaks represent the crystallization of chlorargyrite with pure nanoparticles. It is believed that Ag^+^ from the AgNO3 initially reacted with the chloride ions (Cl^-)^ present in the leaf extract to form chlorargyrite (AgCl). Subsequently, the phytochemicals, especially the phenolic groups of the polyphenols, removed the Ag^+^ from the AgCl to generate pure Ag^0^ crystals (35). The average crystallite size calculated using the Debye−Scherrer equation was 73.67 nm, and the average lattice constant, 0.4089, is quite near to the typical lattice parameter of AgNPs, which is about 0.4086 nm (JCPDS file number 04-0783).

Size distribution profiling by DLS-Zeta technique suggests that these AI-AgNPs are slightly polydisperse with a PDI value of 0.368 (36). The average size of the particles revealed by the intensity vs. size distribution curve was 73.74 nm. The size of the crystallites determined by the Debye-Scherrer equation and the size derived from DLS analysis coincide. The zeta potential of green nanoparticles was -28.04 mV, indicating moderate stability. The particles do not tend to agglomerate, and instead, they have a tendency to repel one another at this large negative zeta potential.

The SEM analysis data reveal the average existence of smooth-surfaced particles with a size of 72.01 nm. This result also supports the results of XRD and DLS. EDAX plot confirms the presence of elemental silver. It also confirms the capping of AgNPs with biogenic materials by showing the presence of carbon and oxygen (37). Chlorine is also indicated by the EDAX plot. Since the proportion of chlorine is far lower than that of silver, this supports the presence of AgCl with AgNPs in a lesser amount.

The TEM investigation provides high-magnification images of AgNPs with an average size of 73.51 nm. The TEM analysis results are also in line with those of XRD, DLS, and SEM.

Infection is the most frequent adverse impact and reason for patient mortality in hospitals. AgNPs-coated plastic catheters exhibit strong in vitro antibacterial activity and inhibit the formation of biofilms from *Enterococcus, Pseudomonas aeruginosa, Candida albicans, Staphylococci*, and *E. Coli* (29). Good antibacterial activity was demonstrated by neem-synthesized AgNPs in our investigation. AI-AgNPs, at an average concentration of 9.4993±0.72679 µg/ml (Mean±SE) (95% CI: 8.0547-10.9439), suppressed the growth of ESBL and MDR *E. coli* and had the ability to kill the bacteria at about 121.0227±7.64143 µg/ml (Mean±SE) (95% CI: 105.8345-136.2109). Earlier research against a variety of microorganisms has also confirmed AI-AgNPs’ antibacterial efficacy (15, 38).

The cytotoxicity of synthesized nanoparticles was evaluated using an MTT assay on HEK293 cell lines. The analysis revealed an IC50 value of 369 µg/ml. A dose-dependent reduction in cell viability was observed with increasing concentrations of AI-AgNPs. Similar findings were reported by Bigdeli et al.(39). Notably, the IC50 value of AI-AgNPs is significantly higher than their antimicrobial concentration, suggesting minimal cytotoxicity combined with strong antimicrobial efficacy.

The present study is limited to in vitro experiments, and cytotoxicity was evaluated using only a single normal cell line, which may not fully represent the complexity of biological systems. Future studies should focus on in vivo infection models to evaluate pharmacokinetics, biodistribution and therapeutic efficacy. If proven safe and effective, neem-mediated AgNPs may have potential applications in antimicrobial coatings, wound dressings, and catheter-associated infection control against drug-resistant pathogens.

## 4. Conclusion

Eradication of infection of ESBL and MDR *E. coli* still causes medical personnel to frown. This study evaluated the efficacy of AgNPs coated with phytochemicals of neem leaf extract in treating AMR *E. coli*. We found that the synthesized nanoparticles are cubic crystals of around 74 nm. Phytochemicals present in the aqueous extract of *A. indica* play a major role in reducing, capping, and stabilizing the nanoparticles. Green synthesized AgNPs showed a significant in vitro antimicrobial activity against ESBL and MDR *E. coli* as demonstrated by the micro-dilution method with minimal cytotoxicity. Therefore, AgNPs synthesized using *A. indica* leaf extract can be utilized as an antimicrobial agent to treat resistant microbial infections. However, an in vivo study is essential to validate this. This eco-friendly method was found to be potent in large-scale, cost-effective production of AgNPs for its use in various medical and biotechnological fields.

## References

1. Mody VV, Siwale R, Singh A, Mody HR. Introduction to metallic nanoparticles. J Pharm Bioallied Sci. 2010;2(4):282–9.

2. Roy P, Das B, Mohanty A, Mohapatra S. Green synthesis of silver nanoparticles using Azadirachta indica leaf extract and its antimicrobial study. Applied Nanoscience. 2017;7(8):843–50.

3. Moore MN. Do nanoparticles present ecotoxicological risks for the health of the aquatic environment? Environment International. 2006;32(8):967–76.

4. Sergeev GB, Shabatina TI. Cryochemistry of nanometals. Colloids and Surfaces A: Physicochemical and Engineering Aspects. 2008;313-314:18–22.

5. Bagur H, Poojari CC, Melappa G, Rangappa R, Chandrasekhar N, Somu P. Biogenically synthesized silver nanoparticles using endophyte fungal extract of Ocimum tenuiflorum and evaluation of biomedical properties. Journal of Cluster Science. 2020;31:1241–55.

6. Ashmore D, Chaudhari A, Barlow B, Barlow B, Harper T, Vig K, et al. Evaluation of E. coli inhibition by plain and polymer-coated silver nanoparticles. Rev Inst Med Trop Sao Paulo. 2018;60:e18.

7. Horie M, Kato H, Endoh S, Fujita K, Komaba LK, Nishio K, et al. Cellular effects of industrial metal nanoparticles and hydrophilic carbon black dispersion. J Toxicol Sci. 2014;39(6):897–907.

8. Bryaskova R, Pencheva D, Nikolov S, Kantardjiev T. Synthesis and comparative study on the antimicrobial activity of hybrid materials based on silver nanoparticles (AgNps) stabilized by polyvinylpyrrolidone (PVP). J Chem Biol. 2011;4(4):185–91.

9. Nymark P, Catalán J, Suhonen S, Järventaus H, Birkedal R, Clausen PA, et al. Genotoxicity of polyvinylpyrrolidone-coated silver nanoparticles in BEAS 2B cells. Toxicology. 2013;313(1):38–48.

10. Franci G, Falanga A, Galdiero S, Palomba L, Rai M, Morelli G, et al. Silver nanoparticles as potential antibacterial agents. Molecules. 2015;20(5):8856–74.

11. Verma A, Mehata MS. Controllable synthesis of silver nanoparticles using Neem leaves and their antimicrobial activity. Journal of Radiation Research and Applied Sciences. 2016;9(1):109–15.

12. Saha J, Begum A, Mukherjee A, Kumar S. A novel green synthesis of silver nanoparticles and their catalytic action in reduction of Methylene Blue dye. Sustainable Environment Research. 2017;27(5):245–50.

13. Ahmed S, Ahmad M, Swami BL, Ikram S. A review on plants extract mediated synthesis of silver nanoparticles for antimicrobial applications: A green expertise. Journal of Advanced Research. 2016;7(1):17–28.

14. Nguyen KC, Seligy VL, Massarsky A, Moon TW, Rippstein P, Tan J, et al., editors. Comparison of toxicity of uncoated and coated silver nanoparticles. Journal of Physics Conference Series; 2013 April 01, 2013.

15. Bhat RS, Almusallam J, Al Daihan S, Al-Dbass A. Biosynthesis of silver nanoparticles using Azadirachta indica leaves: characterisation and impact on Staphylococcus aureus growth and glutathione-S-transferase activity: IET Nanobiotechnol. 2019 Apr 17;13(5):498–502. doi: 10.1049/iet-nbt.2018.5133. eCollection 2019 Jul.

16. Han JW, Gurunathan S, Jeong JK, Choi YJ, Kwon DN, Park JK, et al. Oxidative stress mediated cytotoxicity of biologically synthesized silver nanoparticles in human lung epithelial adenocarcinoma cell line. Nanoscale Res Lett. 2014;9(1):459.

17. Castiglioni S, Caspani C, Cazzaniga A, Maier JA. Short- and long-term effects of silver nanoparticles on human microvascular endothelial cells. World J Biol Chem. 2014;5(4):457–64.

18. Ferdous Z, Nemmar A. Health Impact of Silver Nanoparticles: A Review of the Biodistribution and Toxicity Following Various Routes of Exposure. Int J Mol Sci. 2020;21(7).

19. Subapriya R, Nagini S. Medicinal properties of neem leaves: a review. Curr Med Chem Anticancer Agents. 2005;5(2):149–6.

20. Banerjee P, Satapathy M, Mukhopahayay A, Das P. Leaf extract mediated green synthesis of silver nanoparticles from widely available Indian plants: synthesis, characterization, antimicrobial property and toxicity analysis. Bioresources and Bioprocessing. 2014;1(1):3.

21. Narayanan M, Divya S, Natarajan D, Senthil-Nathan S, Kandasamy S, Chinnathambi A, et al. Green synthesis of silver nanoparticles from aqueous extract of Ctenolepis garcini L. and assess their possible biological applications. Process Biochemistry. 2021;107:91–9.

22. Iskandar K, Molinier L, Hallit S, Sartelli M, Hardcastle TC, Haque M, et al. Surveillance of antimicrobial resistance in low- and middle-income countries: a scattered picture. Antimicrob Resist Infect Control. 2021;10(1):63.

23. Halboup A, Al-Khazzan A, Battah M, Areqi A, Khamaj F, Al-Arifi S. Urinary Tract Infections Management in the Developing Countries. In: Al-Worafi YM, editor. Handbook of Medical and Health Sciences in Developing Countries : Education, Practice, and Research. Cham: Springer International Publishing; 2023. p. 1–19.

24. Ibrahim ME, Bilal NE, Hamid ME. Increased multi-drug resistant Escherichia coli from hospitals in Khartoum state, Sudan. Afr Health Sci. 2012;12(3):368–75.

25. Wilson JW, Schurr MJ, LeBlanc CL, Ramamurthy R, Buchanan KL, Nickerson CA. Mechanisms of bacterial pathogenicity. Postgraduate Medical Journal. 2002;78(918):216–24.

26. Ruiz Ld, Domínguez MA, Ruiz N, Viñas M. Relationship between clinical and environmental isolates of Pseudomonas aeruginosa in a hospital setting. Archives of Medical Research. 2004;35(3):251–7.

27. Peterson E, Kaur P. Antibiotic Resistance Mechanisms in Bacteria: Relationships Between Resistance Determinants of Antibiotic Producers, Environmental Bacteria, and Clinical Pathogens. Front Microbiol. 2018;Volume 9 - 2018.

28. Khalil MMH, Ismail EH, El-Baghdady KZ, Mohamed D. Green synthesis of silver nanoparticles using olive leaf extract and its antibacterial activity. Arabian Journal of Chemistry. 2014;7(6):1131–9.

29. Murei A, Pillay K, Govender P, Thovhogi N, Gitari WM, Samie A. Synthesis, Characterization and In Vitro Antibacterial Evaluation of Pyrenacantha grandiflora Conjugated Silver Nanoparticles. Nanomaterials (Basel). 2021;11(6).

30. Amendola V, Bakr OM, Stellacci F. A Study of the Surface Plasmon Resonance of Silver Nanoparticles by the Discrete Dipole Approximation Method: Effect of Shape, Size, Structure, and Assembly. Plasmonics. 2010;5(1):85–97.

31. Clarkson JP, Winans J, Fauchet PM. On the scaling behavior of dipole and quadrupole modes in coupled plasmonic nanoparticle pairs. Opt Mater Express. 2011;1(5):970–9.

32. Bawoke M. The Impact of Size on the Optical Properties of Silver Nanoparticles Based on Dielectric Function. In: Dr. Alberto J-S, Dr. Gonzalo S, editors. Nanomaterials and Nanostructures - Annual Volume 2024. Rijeka: IntechOpen; 2023. p. Ch. 4.

33. Shaikh WA, Chakraborty S, Owens G, Islam RU. A review of the phytochemical mediated synthesis of AgNP (silver nanoparticle): the wonder particle of the past decade. Applied Nanoscience. 2021;11(11):2625–60.

34. Shankar SS, Rai A, Ahmad A, Sastry M. Rapid synthesis of Au, Ag, and bimetallic Au core–Ag shell nanoparticles using Neem (Azadirachta indica) leaf broth. Journal of Colloid and Interface Science. 2004;275(2):496–502.

35. Hlapisi N, Songca SP, Ajibade PA. Morphological and structural properties of silver/chlorargyrite nanoparticles prepared using Senecio madagascariensis leaf extract and interaction studies with bovine serum albumin. MRS Advances. 2024.

36. Worldwide MI. Dynamic light scattering, common terms defined. Inform white paper Malvern Instruments Limited. 2011;2011:1–6.

37. Ali MH, Azad MAK, Khan KA, Rahman MO, Chakma U, Kumer A. Analysis of Crystallographic Structures and Properties of Silver Nanoparticles Synthesized Using PKL Extract and Nanoscale Characterization Techniques. ACS Omega. 2023;8(31):28133–42.

38. Pachiyappan S, Rashmitha S, Veera P, Ignatious C, Saipriya C, Samrot A. Antibacterial Activity of Neem Extract and its Green Synthesized Silver Nanoparticles against Pseudomonas aeruginosa. Journal of Pure and Applied Microbiology. 2018;12.

39. Bigdeli R, Shahnazari M, Panahnejad E, Cohan RA, Dashbolaghi A, Asgary V. Cytotoxic and apoptotic properties of silver chloride nanoparticles synthesized using Escherichia coli cell-free supernatant on human breast cancer MCF 7 cell line. Artif Cells Nanomed Biotechnol. 2019;47(1):1603–9.

